# High-fat and high-carbohydrate diets worsen the mouse brain susceptibility to damage produced by enterohemorrhagic *Escherichia coli* Shiga toxin 2

**DOI:** 10.1101/2024.02.06.579171

**Authors:** D Arenas-Mosquera, N Cerny, A Cangelosi, PA Geoghegan, EL Malchiodi, M De Marzi, A Pinto, J Goldstein

## Abstract

**Background:** Nutrition quality could be one of the reasons why, in the face of a Shiga toxin-producing enterohaemorrhagic *Escherichia coli* outbreak, some patients experience more profound deleterious effects than others, including unfortunate deaths. Thus, the aim of this study was to determine whether high-fat and/or high-carbohydrate diets could negatively modulate the deleterious action of Shiga toxin 2 on ventral anterior and ventral lateral thalamic nuclei and the internal capsule, the neurological centers responsible for motor activity.

**Methods:** Mice were fed a regular, high-fat, high-carbohydrate diet or a combination of both previous to the intravenous administration of Shiga toxin 2 or vehicle. Four days after intravenous administration, mice were subjected to behavioral tests and then sacrificed for histological and immunofluorescence assays to determine alterations in the neurovascular unit at the cellular and functional levels. Statistical analysis was performed using one-way analysis of variance followed by Bonferroni *post hoc* test. The criterion for significance was p = 0.0001 for all experiments.

**Results:** The high-fat and the high-carbohydrate diets significantly heightened the deleterious effect of Stx2, while the combination of both diets yielded the worst results, including endothelial glycocalyx and oligodendrocyte alterations, astrocyte and microglial reactivity, neurodegeneration, and motor and sensitivity impairment.

**Conclusions:** In view of the results presented here, poor nutrition could negatively influence patients affected by Stx2 at a neurological level. Systemic effects, however, cannot be ruled out.

## Background

Shiga toxin (Stx) from Stx-producing *Escherichia coli* (STEC) is the virulence factor responsible for hemorrhagic colitis, hemolytic uremic syndrome (HUS), and associated acute encephalopathy, triggered by the presence of STEC in food, water, or cross-contamination. When the central nervous system (CNS) is compromised, mortality rates increase significantly (1).

Stx2 is encoded by a lambda bacteriophage. Through the activation of the SOS response in bacteria, the *Stx2* gene undergoes excision and replication, and Stx2 is then expressed and released (2–4). It is well accepted that Stx2 produces its deleterious effect via its canonical cell membrane receptor globotriaosylceramide (Gb3) (5). Our group and others have reported the localization of this receptor in neurons and microglia in the CNS (1, 5, 6). In addition, Gb3 is upregulated following Stx2 administration, concomitantly with a wide range of neurological alterations (5, 7–9). Acute encephalopathy is a sign of poor prognosis which increases mortality rates in children suffering from HUS (10). Neurological damage signs and symptoms include decerebrate posture, hemiparesis, ataxia and cranial nerve palsy, hallucinations, seizures, and changes in the level of consciousness (from lethargy to coma) (11, 12). Similar signs of neurological impairment observed in mouse models of HUS encephalopathy include lethargy, shivering, abnormal gait, hindlimb paralysis, spasm-like seizures, reduced spontaneous motor activity, and pelvic elevation (6, 13).

The thalamus is among the brain areas most affected in patients with HUS encephalopathy (8, 14–16). Patients usually exhibit symmetrical hyper-intensities of thalamic magnetic resonance imaging, which is consistent with thalamic damage (14–16). Neurological signs comprise alterations in consciousness, cognitive dysfunction Lattention, orientation, working or short-term memory deficitsL, and apraxia. Additional clinical signs include headaches and focal neurological deficits such as extra pyramidal, cerebellar, or brainstem symptoms (8). As the thalamus is a diencephalic relay structure for major ascending and descending pathways, all these signs may reflect thalamic damage.

Strikingly, it is not clear why only 20% of children infected with STEC develop HUS, with 5% of these children also developing HUS-associated encephalopathy (1). A plausible explanation may lie in poor nutrition based on a high-fat and/or high-carbohydrate diet, consistent with a chronic systemic pro-inflammatory state and increased oxidative stress (17). We therefore hypothesize that the quality of a high-fat and/or high-carbohydrate diet in children suffering STEC may determine the severity of HUS encephalopathy. For these reasons, the aim of this study is to determine whether high-fat and/or high-carbohydrate diets modulate the deleterious action of Stx2 on the thalamic neurovascular unit responsible for motor activity in mice, particularly ventral anterior and ventral lateral nuclei (VA-VL) and the internal capsule (18), an ascending pathway connecting fibers from the thalamus with the cerebral cortex (19). This work also aims to establish whether a pro-inflammatory component is involved in Stx2 effects and whether the changes observed at a cellular level are associated with mouse behavior. This study may thus help understand the pathophysiological events occurring in a translational murine model of STEC intoxication which attempts to reproduce clinical events from a dietary perspective.

## Methods

### Animals

Male NIH Swiss mice were housed under 12h-light/12h-dark conditions. Food and water were provided *ad libitum* (as described below), and the experimental protocols and euthanasia procedures were reviewed and approved by the Institutional Committee for the Care and Use of Laboratory Animals of the School of Medicine, Universidad de Buenos Aires, Argentina (Resolution N° 3020/2019). All the procedures were performed in accordance with the EEC guidelines for the care and use of experimental animals (EEC Council 86/609).

### Diets

Twenty one-day-old weaning mice were divided into four groups according to their diet composition (Asociación de CooperativasArgentinas, San Nicolás, Buenos Aires, Argentina) as follows (Table 1): control normal diet (ND); high-fat diet (HF) consisting of a ND enriched with an additional 37.97% fat; high-carbohydrate (HC) consisting of a ND enriched with 10% saccharose solution (#9789.08, Biopack, Buenos Aires, Argentina); and a combined HF+HC diet. After the weaning day, the four diets were administered for 25 days until the end of the protocol. On day 21 (experimental day zero), animals were subjected to intravenous (i.v.) administration (100µl) of either Stx2 (1ng) or vehicle (saline solution). The Stx2 dose was about 60% of the LD_50_ (1.6ng per mice). Thus, each diet group (ND, HF, HC and HF+HC) was in turn divided into two i.v. treatments (+veh or +Stx2, n=10). Four days after i.v. treatment, animals were subjected to behavioral tests and then sacrificed for immunofluorescence assays.

**Table 1:** Nutritional composition for the four groups (%).

| Source | ND | HF | HC | HF+HC |
| --- | --- | --- | --- | --- |
| Protein | 32.57 | 17.93 | 32.57 | 17.93 |
| Carbohydrate | 60.49 | 37.59 | 60.49 | 37.59 |
| Fat | 6.50 | 44.47 | 6.50 | 44.47 |
| --- | --- | --- | --- | --- |
| Saccharose | ----- |  | 10.00 | 10.00 |

### Swimming test

A swimming test apparatus was built to evaluate mouse motor behavior. The apparatus consisted of a glass tank of 100-cm length, 6-cm width, and 30-cm depth, filled with water (23 °C) up to 10 cm from the top. A visible escape platform was placed 0.5cm above water at the end of the apparatus. On the opposite side of the escape platform, a vertical dotted black line was drawn 60cm from the platform. This line served as the start line for recording swimming performance. For this purpose, each mouse was placed before the vertical dotted black line and learned to swim straight into the visible escape platform on the opposite side after two trials. Mouse performance was recorded, and the latency to swim the 60-cm distance was registered.

### Von Frey test

To determine mouse sensitivity, paw withdrawal to mechanical stimuli was measured using the Von Frey test. The test consisted of a 0.7-cm x 0.7-cm chamber in which mice were placed on top of a wire rack with a 10-cm x 12.5-cm grid and allowed to move freely. Animals were habituated to the chamber for 5 minutes before mechanical stimuli were applied.

The hind paw withdrawal threshold was determined using a sequence of ascending mechanical stimuli (expressed in grams) of calibrated von Frey monofilaments. Twenty filaments ranging from 1.65 g to 6.65 g were used by experimenters who were blinded to mouse treatment allocation. Stimuli were applied in the central region of the plantar surface, avoiding the foot pads. The filament was applied only when the mouse was stationary and standing on all four paws. The withdrawal response was considered valid only if the hind paw was completely removed from the platform. The threshold was determined on the right hind limb.

### Treatments and sample processing

After swimming and Von Frey tests, animals were sacrificed, and brains were processed for histological and immunofluorescence assays. Briefly, mice were anesthetized with pentobarbital (100mg/kg) and intracardially perfused with paraformaldehyde 4% in phosphate buffer saline (PBS) 0.1M (pH 7.4) at 4 °C. Brains were removed from the skull, post-fixed with the same solution overnight, and cryopreserved with sucrose solutions at three different concentrations (10, 20 and 30%) overnight. Coronal sections of 20 µm were obtained on a cryostat and stored in a cryopreserved solution (30% ethylene glycol and 20% glycerol in PBS 0.1M) at −20 °C until the day of the assays.

### Histological and immunofluorescence assays

To determine the effect of Stx2 on vascular endothelial cells, brain slices were incubated with biotinylated lectin (10μg/ml in PBS 0.1M with 0.3% Triton X-100; Sigma, St. Louis, MO, USA) for 24 hours at 4 °C and subsequently incubated with streptavidin Alexa-488 (Invitrogen Molecular Probes, Carlsbad, CA, USA) in a 1:500 dilution for 1 hour at room temperature.

To determine the effect of Stx2 on neurons, astrocytes, microglia and oligodendrocytes, brain slices were incubated with mouse neuronal nuclei antibody (anti-NeuN, 1:500; Millipore, Temecula, CA, USA), rabbit glial fibrillary acidic protein antibody (anti-GFAP,1:500; Dako, Glostrup, Denmark), goat ionized calcium binding adaptor molecule 1 antibody (anti-Iba1,1:500; Millipore), and mouse myelin basic protein antibody (anti-MBP, 1:500, Dako, Glostrup, Denmark), respectively for 24 hours at 4 °C. After several washes, slices were incubated with their respective secondary antibodies: goat anti-mouse Alexa Fluor 555 (1:500; Amersham, GE, Piscataway, NJ, USA), goat IgG anti-rabbit Alexa Fluor 555 (Invitrogen Molecular Probes, Carlsbad, CA, USA), and donkey anti-goat Alexa Fluor 488 (1:500; Millipore). Hoechst 33342 (1:500; Sigma) was used in all histological and immunofluorescence assays to detect cell nuclei.

The areas chosen in histological and immunofluorescence sections were observed with an Olympus BX50 epifluorescence microscope provided with a Cool-Snap digital camera and a confocal Olympus FV1000 microscope.

### Image analysis

All micrographs were analyzed using FIJI ImageJ software. For lectin, a scale was first set up using a Neubauer chamber. Next, 8-bit newly transformed micrographs were adjusted with the “threshold” tool and then analyzed with the “set measurements” tool. The number and percentage of lectin-positive particles were then determined. The “cell counter” plugin was used to establish the percentage of morphologically damaged nuclei-neurons and the number of Iba1-positive cells. The “ROI manager” tool was used to establish the expression levels of GFAP, MBP, and Iba1.

### Statistical analysis

The data are presented as the mean ± SEM. In all assays, statistical analyses were conducted using one-way analysis of variance (ANOVA) followed by Bonferroni *post hoc* test (GraphPad Prism 8, GraphPad Software, Inc., San Diego, CA, USA) between i.v. treatments (control and Stx2). The criterion for significance was p<0.0001 for all experiments.

## Results

### The HF and/or HC diets heightened Stx2-induced alterations in the microvessel profile

The area observed in this study is located along the VA and VL thalamus (Fig. 1A). Lectins from *Lycopersicum esculentum,* a marker that binds to the glycocalix of microvessels, was used to assess morphological changes promoted by Stx2 in each diet (Fig. 1). Parameters evaluated through lectin histofluorescence included the number of lectin-positive particles (Fig. 1K), microvessel size (Fig. 1L), and percentage of area occupied by microvessels (Fig. 1M).

**Figure 1.**
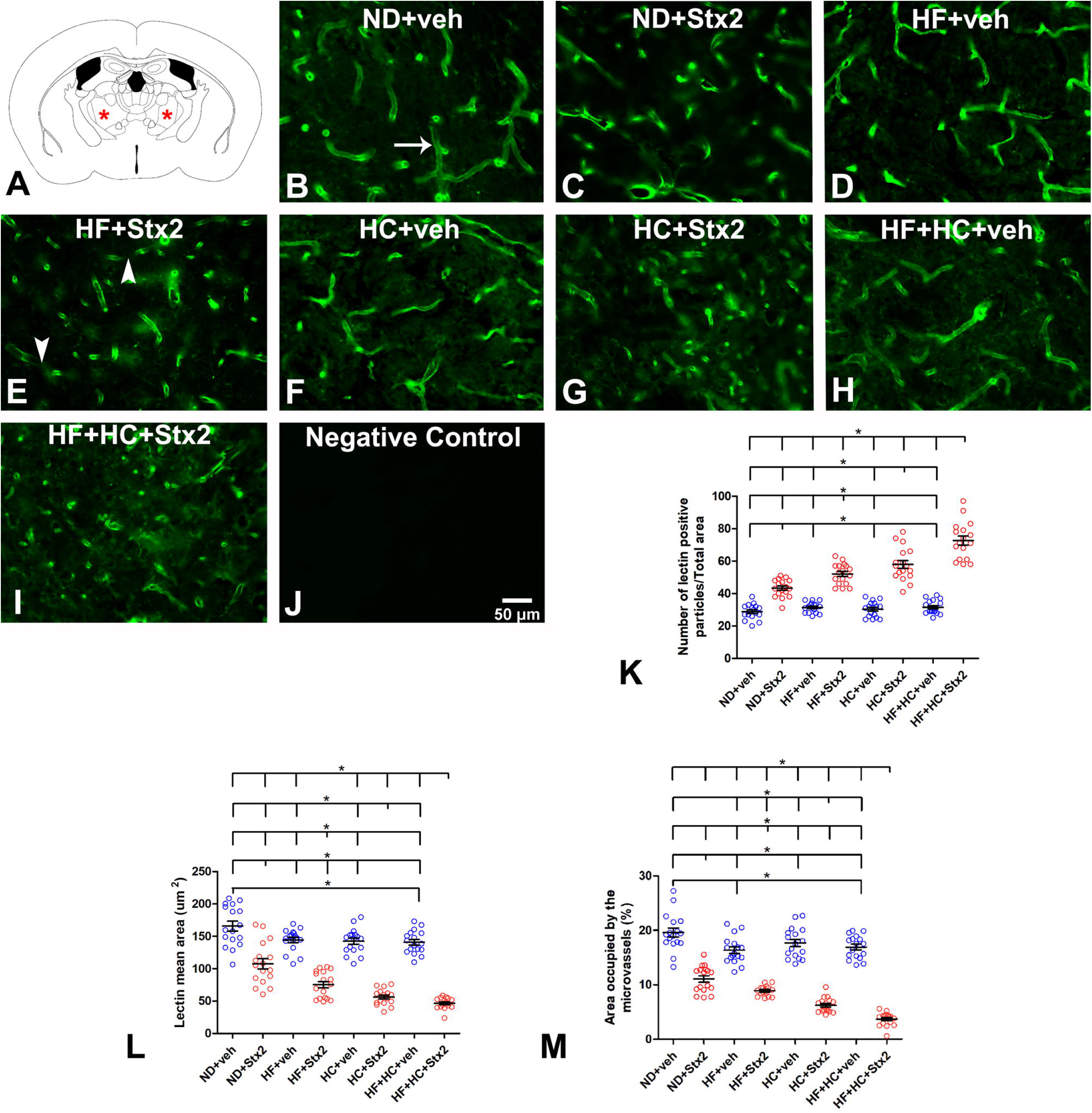
Changes in microvasculature lectin fluorescence pattern. A: the murine VA-VL brain area studied; B: micrograph shows the microvasculature profile in the ND+veh control group; C: ND+Stx2 treatment; D: HF+veh treatment; E: HF+Stx2 treatment; F: HC+veh treatment; G: HC+Sxt2 treatment; H: HF+HC+veh; I: HF+HC+Stx2; J: negative control by not adding Lycopersicum esculentum lectins; K: number of lectin-immunopositive particles; L: size of microvessels (µm2); M: area occupied by microvessels (%); arrow in B: a conserve microvessel; arrowheads in E: fragmented microvessels. Data were analyzed by one-way ANOVA and Bonferroni post hoc test, p = 0.0001, n=5. Scale bar in J applies to all panels.

#### Microvessel assessment in diet groups treated with vehicle

The ND+veh thalamus showed a conserved pattern and continuous lectin histofluorescence binding throughout microvessels belonging to endothelial cell membranes (Fig. 1B, arrow). The number of microvessels bound to lectins-(table 2) observed as histofluorescent lectin-positive particlesL showed no significant differences across veh-treated diet groups (Fig. 1K), as seen in panels of Fig. 1 B, D, F, H.

**Table 2:** number of microvessels bound to lectins; p = 0.0001.

| ND+veh | ND+Stx2 | HF+veh | HF+Stx2 |
| --- | --- | --- | --- |
| 28.82±4.45 | 43.29±5.24 | 31.41±3.11 | 52.06±6.24 |
| HC+veh | HC+Stx2 | HF+HC+veh | HF+HC+Stx2 |
| 30.29±4.57 | 57.94±9.89 | 31.53±4.12 | 72.65±11 |

However, microvessel size (Table 3) was significantly smaller in HF+HC+veh (Fig. 1H) than in ND+veh mice (Fig. 1B), with no significant differences between ND+veh (Fig. 1B), HF+veh (Fig. 1D), or HC+veh (Fig. 1H) mice, Fig. 1L.

**Table 3:** Microvessel size (um^2^); p = 0.0001.

| ND+veh | ND+Stx2 | HF+veh | HF+Stx2 |
| --- | --- | --- | --- |
| 165.86±30.32 | 107.63±31.58 | 145.27±17.30 | 81.27±18.71 |
| HC+veh | HC+Stx2 | HF+HC+veh | HF+HC+Stx2 |
| 142.63±18.87 | 57.27±11.99 | 139.27±19.14 | 46.57±8.23 |

Finally, the percentage of area occupied by microvessels was significantly smaller in HF+veh (Fig. 1D) and HF+HC+veh (Fig. 1H), as compared to ND+veh mice (Fig. 1B) (Table 4, Fig. 1M). Surprisingly, HF and HF+HC diets reduced the area occupied by microvessels regardless of Stx2 administration.

**Table 4:** total area occupied by microvessels (%); p = 0.0001.

| ND+veh | ND+Stx2 | HF+veh | HF+Stx2 |
| --- | --- | --- | --- |
| 18.41±2.53 | 10.31±1.95 | 17.05±2.88 | 8.52±0.78 |
| HC+veh | HC+Stx2 | HF+HC+veh | HF+HC+Stx2 |
| 17.36±2.90 | 5.71±1.00 | 16.77±2.27 | 3.13±1.13 |

#### Microvessel assessment in diet groups treated with Stx2

We have previously reported that the administration of a sublethal dose of Stx2 alters the microvessel profile (8). In the present study, Stx2 administration combined with HF and/or HC diets further worsened the microvessel profile, as observed in HF+veh, HC+veh and HF+HC+veh mice (Fig. 1B-I). Indeed, Stx2 administration combined with the HF+HC diet induced a peak in the number of histofluorescent lectin-positive particles (fragmented particles) (Fig. 1K), followed by HF+Stx2 and HC+Stx2 as compared to their respective controls (Table 2, Fig. 1K).

Stx2-treated microvessels presented non-continuous fluorescence labeling (Fig.1E arrowheads). As a distinct feature of microvessel injury, these non-continuous fluorescence microvessels have been proven smaller in Stx2-treated thalami than in controls (8). In our work, microvessel size reached the lowest value in HF+HC+Stx2 mice, followed by HF+Stx2 and HC+Stx2 (Table 3, Fig. 1L).

Furthermore, the total area occupied by microvessels was significantly smaller in Stx2-treated mice, as observed in other brain areas (8, 9). The HF+HC+Stx2 group rendered the smallest area, followed by HF+Stx2 and HC+Stx2 (Table 4, Fig. 1M). In contrast, no lectin histofluorescence binding was observed in negative controls (Fig. 1J).

### The HF+HC diet magnified Stx2-induced astrocyte reactivity

To determine the effect of HF and/or HC diets on astrocyte reactivity, GFAP expression was assessed in these cells in the internal capsule (Fig. 2A); this region consists of afferent and efferent cortical fibers, the anterior part containing ascending thalamic projections (21). Significant astrocyte reactivity was observed in the four diet groups after Stx2 treatment as compared to their respective veh-treated groups (Table 5, Fig. 2B-K). Maximal astrocyte reactivity was found in the HF+HC+Stx2 group, with no significant differences across the other Stx2-treated groups (Fig. 2C, E, G, I). Finally, no specific immunofluorescence for GFAP expression was observed in negative controls (Fig. 2J).

**Figure 2.**
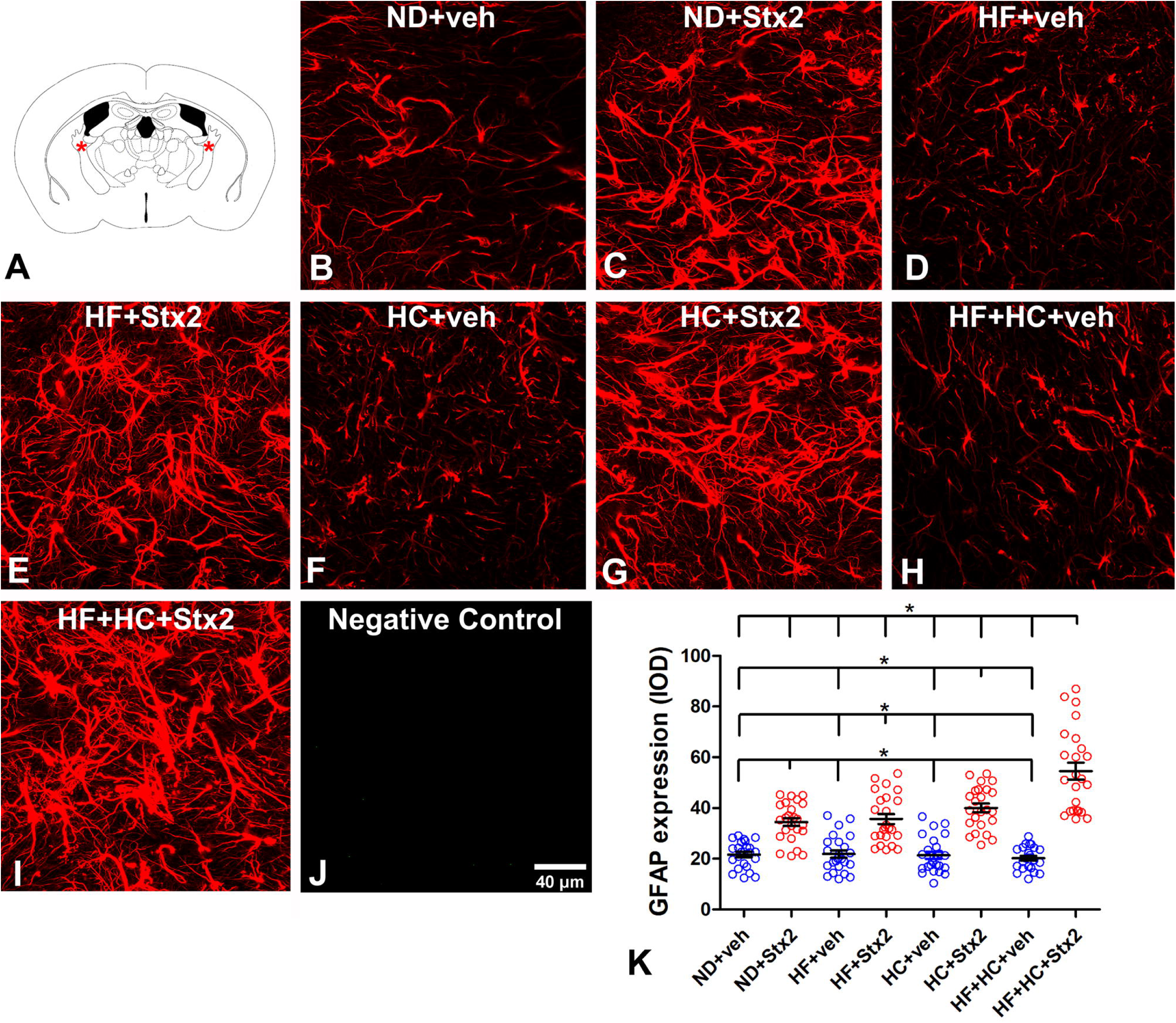
Changes in astrocyte GFAP expression. A: the murine internal capsule brain area studied; B: micrograph shows the astrocyte expression of GFAP in the ND+veh control group; C: ND+Stx2 treatment; D: HF+veh treatment; E: HF+Stx2 treatment; F: HC+veh treatment; G: HC+Stx2 treatment; H: HF+HC+veh treatment; I: HF+HC+Stx2 treatment; J: negative control by omitting primary antibody; K: quantification of GFAP expression levels in all treatments. Data were analyzed by one-way ANOVA and Bonferroni post hoc test, p = 0.0001, n=5. Scale bar in J applies to all panels.

**Table 5:** Astrocyte reactivity (IOD); p = 0.0001.

| ND+veh | ND+Stx2 | HF+veh | HF+Stx2 |
| --- | --- | --- | --- |
| 21.65±5.20 | 34.50±7.68 | 21.83±7.20 | 35.68±9.66 |
| HC+veh | HC+Stx2 | HF+HC+veh | HF+HC+Stx2 |
| 21.38±7.07 | 40.08±8.49 | 20.23±4.48 | 54.53±16.42 |

### The HF and/or HC diets worsened Stx2-induced neurodegeneration

NeuN was used as a biomarker to determine neurodegenerative events (22) in VA-VL thalamus (Fig. 3A). Conserved NeuN immunofluorescence was detected in veh-treated neuronal nuclei (Fig. 3B, arrowhead). A non-significant number of abnormal thalamic nuclei was detected in veh-treated groups regardless of diets (Table 5, Fig. 3K), all of them showing conserved NeuN immunofluorescence colocalizing with Hoechst staining (Fig. 3B, D, F, H, K, see arrowhead in B). In contrast, Stx2-treated groups showed significant aberrant neuronal phenotypes (Fig. 3C, E, G, I), a neurodegenerative profile evidenced by NeuN immunofluorescence in the perinuclear zone but not in the nucleus (Fig. 3I, arrow). The number of neurons bearing an aberrant phenotype was maximal in the HF+HC+Stx2 group (Table 6, Fig. 3K). No NeuN immunofluorescence was observed in negative controls (Fig. 3J).

**Figure 3.**
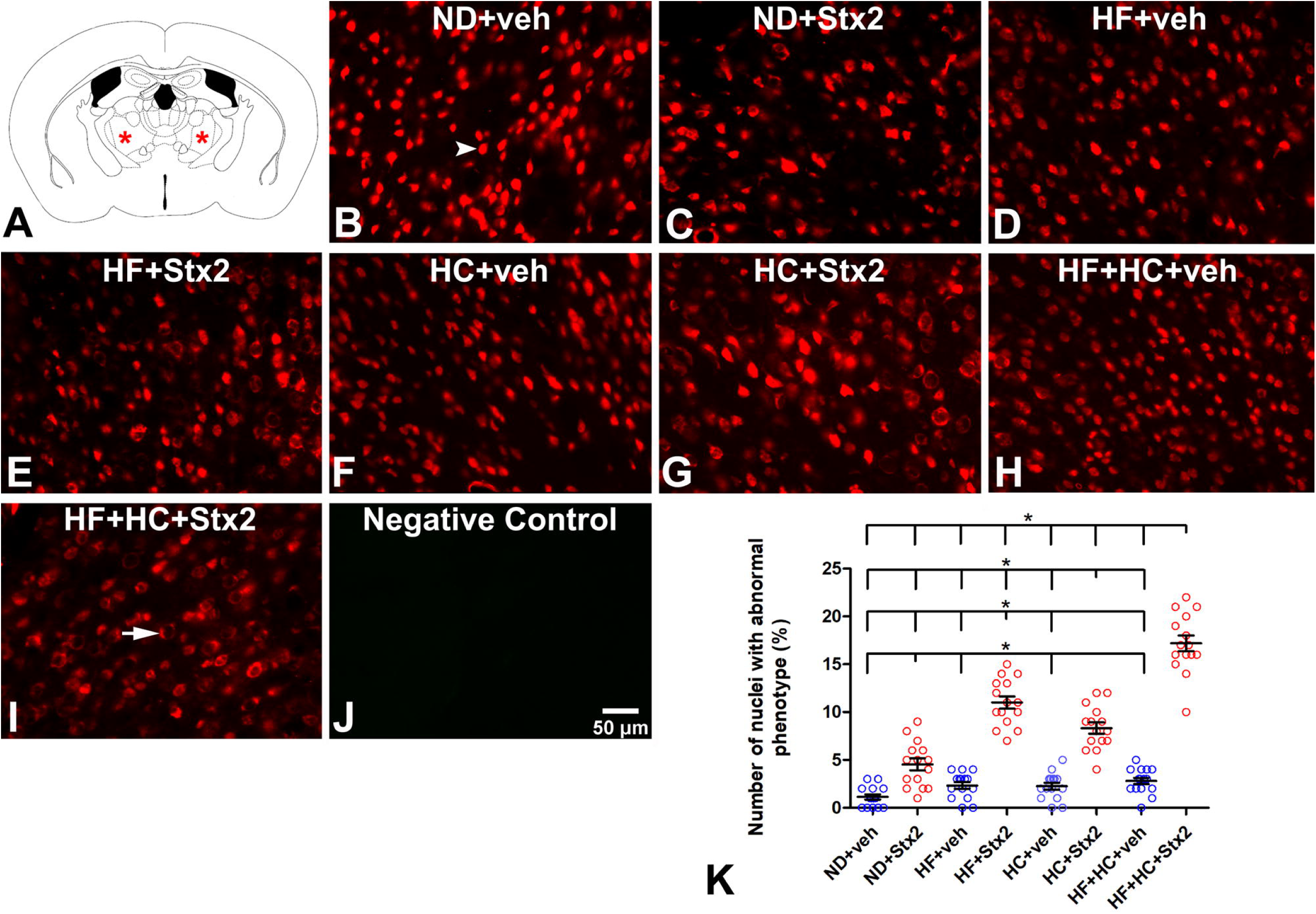
Changes in the neuron NeuN fluorescence patterns. A: the murine VA-VL brain area studied; B: micrograph shows NeuN localization in the ND+veh control group; C: ND+Stx2 treatment; D: HF+veh treatment; E: HF+Stx2 treatment; F: HC+veh treatment; G: HC+Stx2 treatment; H: HF+HC+veh treatment; I: HF+HC+Stx2 treatment; J: negative control by omitting primary antibody; K: quantification of nuclei with abnormal phenotypes in all treatments. Arrowhead in B: normal phenotype; arrow in I: abnormal phenotype. Data were analyzed by one-way ANOVA and Bonferroni post hoc test, p = 0.0001, n=5. Scale bar in J applies to all panels.

**Table 6:** number of nuclei with abnormal phenotype; p = 0.0001.

| ND+veh | ND+Stx2 | HF+veh | HF+Stx2 |
| --- | --- | --- | --- |
| 1.13±1.06 | 4.53±2.39 | 2.33±1.35 | 11.00±2.45 |
| HC+veh | HC+Stx2 | HF+HC+veh | HF+HC+Stx2 |
| 2.27±1.39 | 8.33±2.29 | 2.80±1.32 | 17.20±3.12 |

### The HF+HC diet boosted Stx2-induced microglia reactivity

As Iba1 is a microglia/macrophage-specific calcium-binding protein and a marker of activated microglia (23), Iba1 expression levels were used to assess microglial activation in the VA-VL thalamus (Fig. 4A). A significant rise was observed in Iba1 expression in HC+veh and HF+HC+veh in comparison with ND+veh (Table 7, Fig. 4 B, D, F, H, K). At variance, a significant increase in Iba1 expression levels was found in all four Stx2 groups (Table 7, Fig. 4C, E, G, I, K). The maximal increase in Iba1 expression was observed in the HC+HF+Stx2 condition (Fig.4I, K). No Iba1 immunofluorescence was observed in negative controls omitting the primary antibody (Fig. 4J).

**Figure 4.**
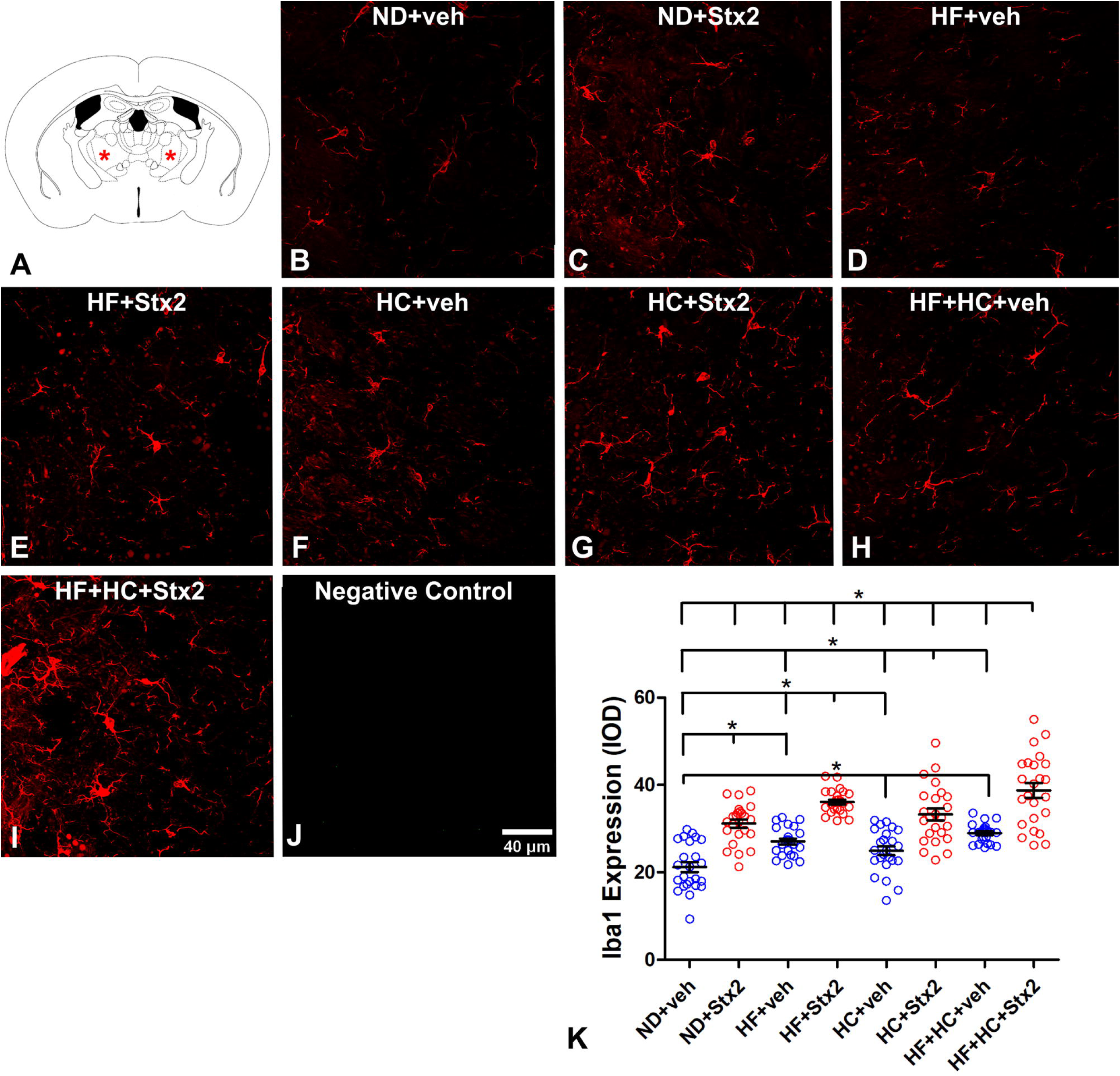
Changes in Iba1 expression and number of Iba1-positive cells. A: the murine VA-VL brain area studied; B: micrograph shows the expression of Iba1 and the number of Iba1-positive cells in the ND+veh control group; C: ND+Stx2 treatment; D: HF+veh treatment; E: HF+Stx2 treatment; F: HC+veh treatment; G: HC+Stx2 treatment; H: HF+HC+veh treatment; I: HF+HC+Stx2 treatment; J: negative control by omitting primary antibody; K: quantification of Iba1 expression levels in all treatments. Data were analyzed by one-way ANOVA and Bonferroni post hoc test, p = 0.0001, n=5. Scale bar in J applies to all panels.

**Table 7:** microglial reactivity (IOD); p = 0.0001.

| ND+veh | ND+Stx2 | HF+veh | HF+Stx2 |
| --- | --- | --- | --- |
| 21.21±5.47 | 31.17±4.53 | 27.05±3.20 | 36.14±2.74 |
| HC+veh | HC+Stx2 | HF+HC+veh | HF+HC+Stx2 |
| 24.91±4.96 | 33.26±6.67 | 28.95±2.13 | 38.75±8.18 |

### The HF+HC diet deepened Stx2-induced reduction in oligodendrocyte MBP expression

MBP is a myelin sheath protein whose loss is considered a marker of myelin degeneration (24). For this reason, changes in MBP expression were assessed in the internal capsule (Fig. 5A) to determine myelin degeneration following diets and treatments. We have previously shown that Stx2 reduces the expression of MBP in mice following a regular diet (20, 25). In this study, no significant differences were observed across veh-treated groups (Table 8, Fig. 5K). However, all Stx2 groups underwent a significant reduction in MBP expression (Fig. 5K), with the minimum value observed in HF+HC+Stx2 animals (Table 8). No MBP immunofluorescence was observed in negative controls (Fig. 5J).

**Figure 5.**
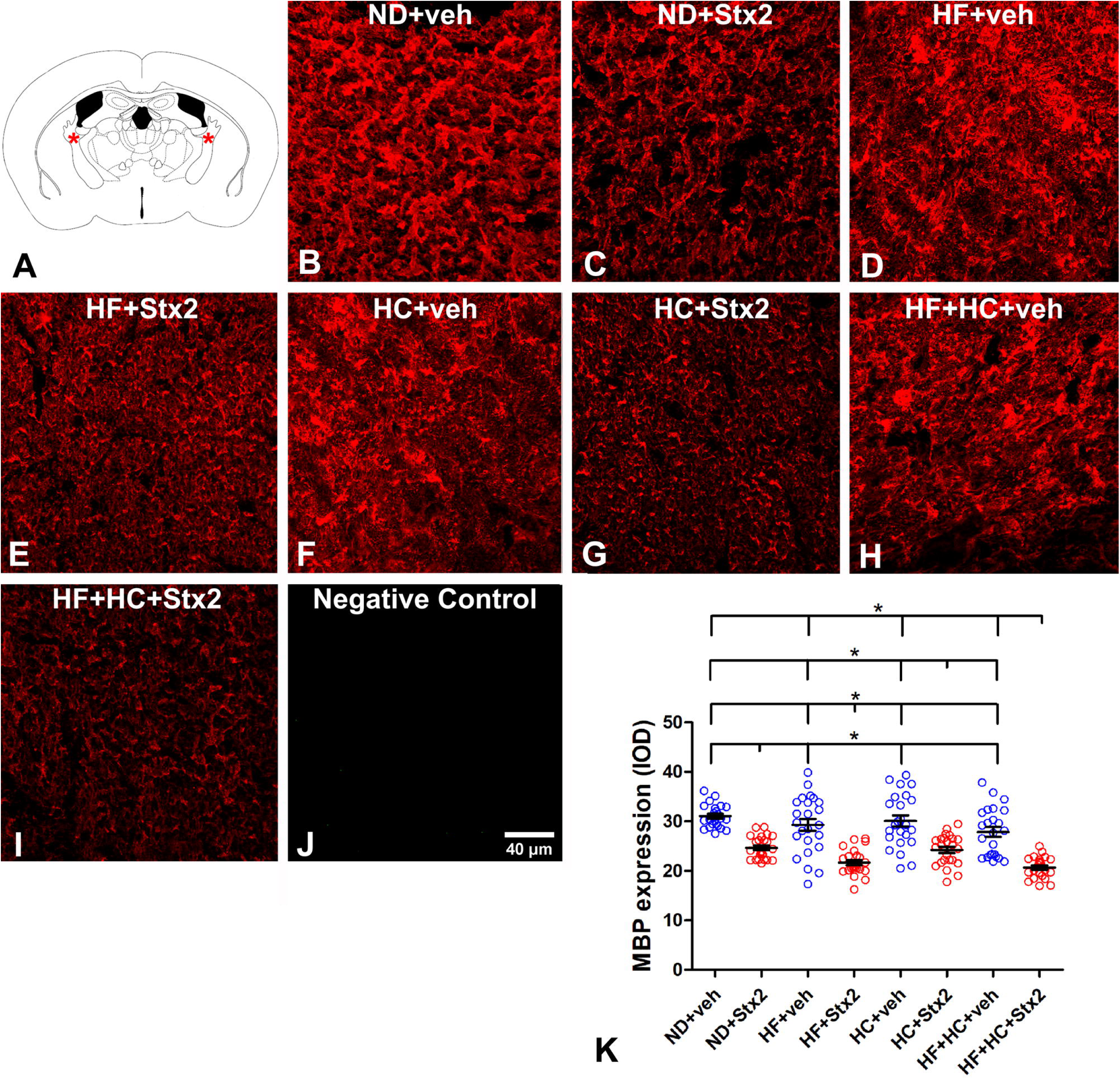
Changes in MBP expression. A: the murine internal capsule brain area studied; B: micrograph shows the expression of MBP in the ND+veh control group; C: ND+Stx2 treatment; D: HF+veh treatment; E: HF+Stx2 treatment; F: HC+veh treatment; G: HC+Stx2 treatment; H: HF+HC+veh treatment; I: HF+HC+Stx2 treatment; J: negative control by omitting primary antibody; K: quantification of MBP expression levels in all treatments. Data were analyzed by one-way ANOVA and Bonferroni post hoc test, p<0.005, n=5. Scale bar in J applies to all panels.

**Table 8:** MBP expression (IOD); p = 0.0001.

|  |  |  |  |
| --- | --- | --- | --- |
| ND+veh | ND+Stx2 | HF+veh | HF+Stx2 |
| 31.05±2.27 | 24.65±2.28 | 29.30±5.87 | 21.68±2.56 |
| HC+veh | HC+Stx2 | HF+HC+veh | HF+HC+Stx2 |
| 30.10±5.44 | 24.23±3.08 | 27.85±4.88 | 20.69±2.09 |

### The HF+HC diet lowered the motor performance in the context of Stx2

Mice were subjected to a swimming motor test (Fig. 6A) to establish whether cellular damage induced by Stx2 and deepened by the HF and/or HC diets correlated with neurological motor alterations (26). All Stx2-treated groups required significantly more time to reach the platform than their respective veh-treated groups (Fig. 6B). No significant differences in latency were found across Stx2-treated groups, although a non-significant peak was observed in the HF+HC+Stx2 group, indicative of motor deficits (Table 9, Fig. 6B).

**Figure 6.**
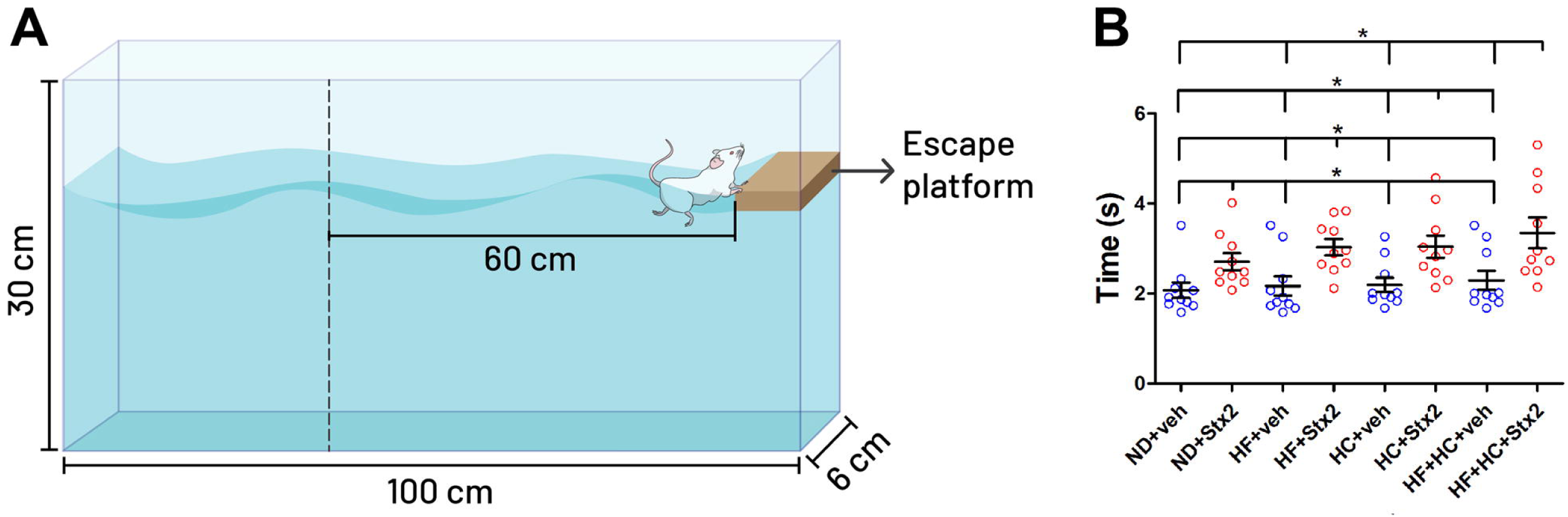
Changes in mouse motor behavior. A: design of the test device; B: columns indicate the different treatments. Data were analyzed by one-way ANOVA and Bonferroni post hoc test, p = 0.0001, n=10.

**Table 9:** latency time (seconds); p = 0.0001.

| ND+veh | ND+Stx2 | HF+veh | HF+Stx2 |
| --- | --- | --- | --- |
| 2.07±0.55 | 2.70±0.60 | 2.17±0.68 | 3.03±0.57 |
| HC+veh | HC+Stx2 | HF+HC+veh | HF+HC+Stx2 |
| 2.19±0.52 | 3.04±0.79 | 2.29±0.67 | 3.35±1.08 |

### Stx2 affected mouse sensitivity without diet involvement

As neurological motor performance was hindered in mice treated with Stx2, we next evaluated mouse sensitivity using Von Frey filaments (Fig. 7A). Changes in paw withdrawal to mechanical stimuli with different filament sizes determine the degree of alterations in sensitivity (27). No significant differences were found in mouse paw sensitivity across Stx2-treated groups (Fig. 7B). Worth pointing out, stronger filaments were used in Stx2-treated animals, which reflects significantly lower sensitivity in Stx2 groups as compared to their respective veh-treated groups (Table 10, Fig. 7B).

**Figure 7.**
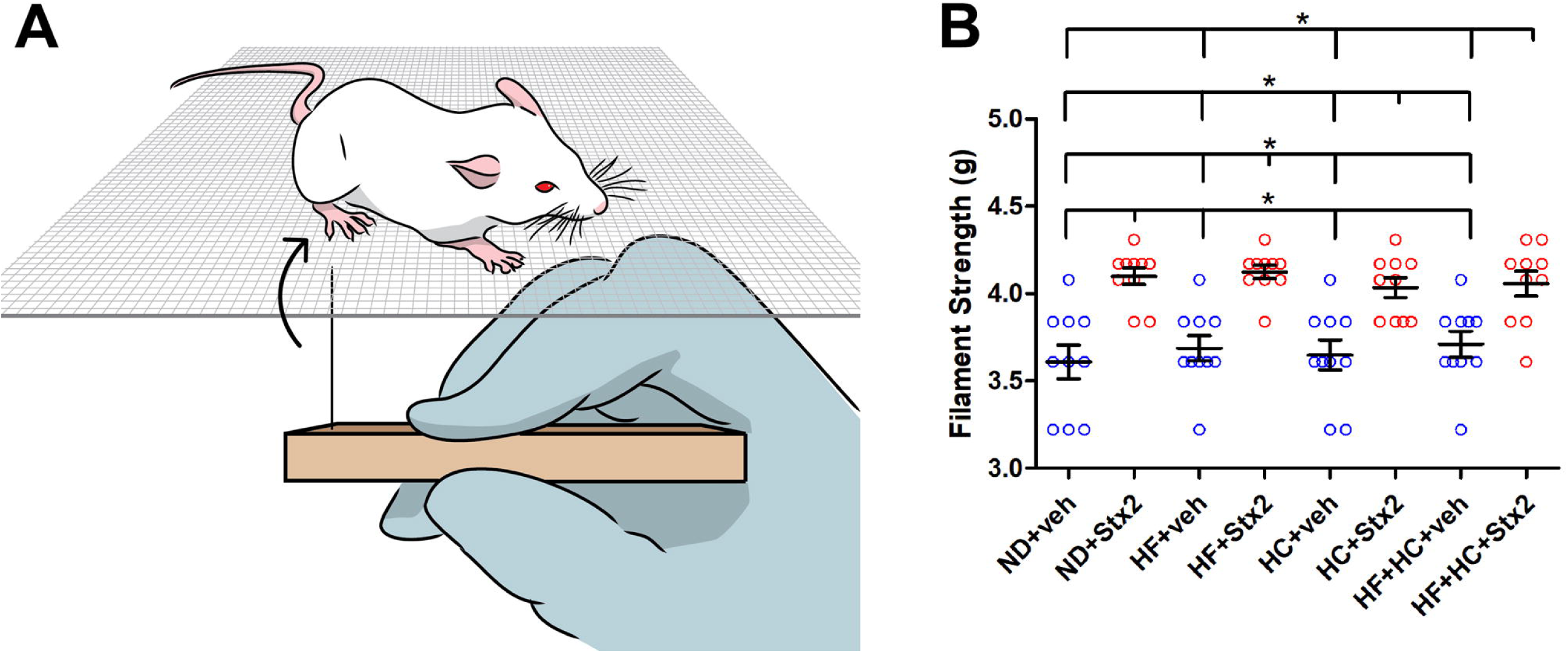
Changes in mouse sensitivity. A: design of the test device; B: columns indicate the different treatments. Data were analyzed by one-way ANOVA and Bonferroni post hoc test, p = 0.0001, n=10.

**Table 10:**
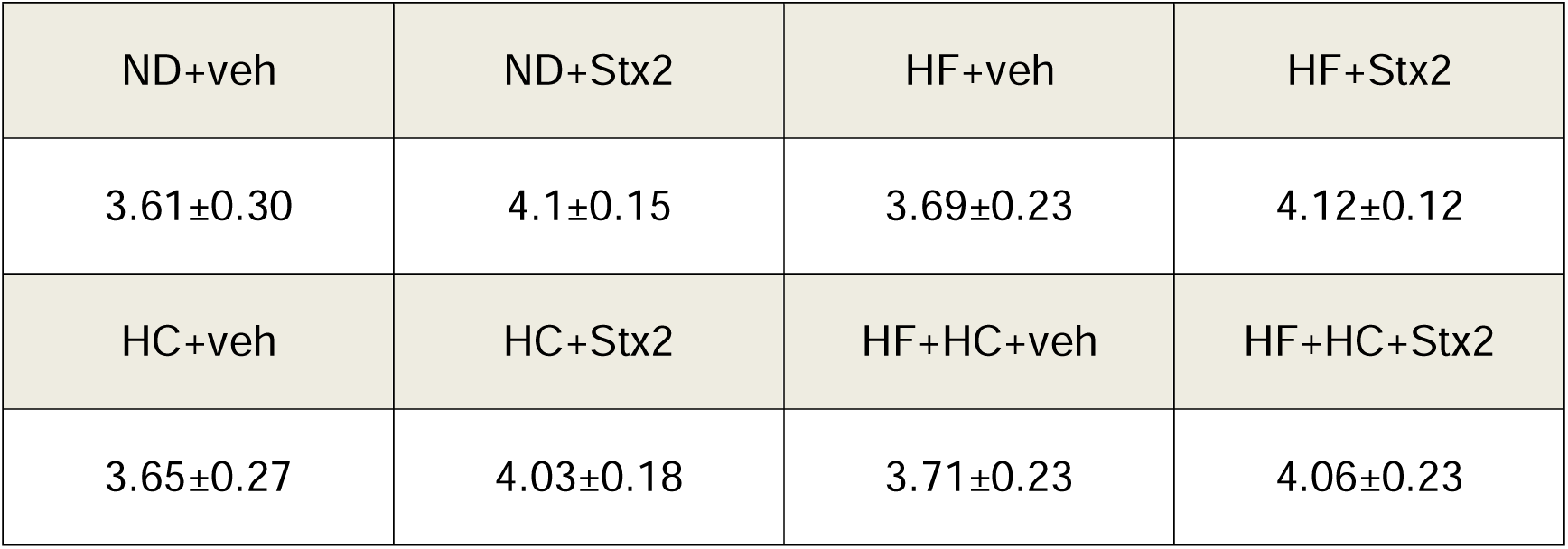
sensitivity (grams); p = 0.0001.

## Discussion

The present findings demonstrate the implications of a poor diet, which may define the health condition of patients intoxicated with STEC. To the best of our knowledge, the results presented here are novel, as they demonstrate the relevance of nutrition in encephalopathy triggered by Stx2 for the first time. These data could explain why, given a STEC outbreak, some patients suffer extreme, potentially lethal neurological alterations, while others present only mild and reversible symptoms (10). Previous reports have demonstrated that high-fat and high-sugar diets promote cognitive and behavioral alterations, together with an increase in systemic and local proinflammatory cytokines (28, 29). In the present study, the HC and HF diets worsened the damage produced by Stx2 in the mouse motor thalamus and internal capsule. These findings were obtained at a cellular level, with microvessel profile alterations and astrocyte and microglial reactivity. Neurodegeneration and myelin protein alterations were concomitantly observed, all indicative of a pathological state. Similar findings on peripheral sensory-motor dysfunctions and neurodegeneration in the context of reactive pro-inflammatory microglia have been reported in brain areas in a model of aging mice subjected to a western diet Lhigh-fat, high-sugar, and high-saltL (30). Similarly, elderly rats subjected to a high-fat, high-sugar diet evidenced impaired spatial learning and working memory and increased anxiety-like behavior, associated with decreased neurogenesis and increased neuroinflammation fostered by astrogliosis (31). Furthermore, a model of metabolic syndrome has revealed a proinflammatory state Lincluding exacerbated oxidative stress and neuronal damageL in reptilian and limbic murine brain regions, resulting in recognition memory loss (32).

Microbiota, the intestine, and the brain interact in a multidirectional way, mainly through the vagus nerve. The afferent fibers of the vagus nerve constitute the main parasympathetic link of the gut-brain axis, as their axonal terminals in the intestinal mucosa sense the mediators released by intestinal epithelial cells, as well as substances released by intestinal bacteria such as LPS (33–35). It has been established that dietary habits influence intestinal microbiome diversity and can induce profound changes in brain homeostasis. These changes may be mediated by microglial activation, systemic inflammation, and vagal afferent signaling, culminating in neuroinflammation. Eating habits have changed significantly in recent years toward a diet rich in processed fats and low in fibers and other functional components of foods. This diet, commonly referred to as western diet, favors the development of metabolic alterations in multiple organs including the CNS (36). The intestine is first damaged, as a western diet alters its ability to function as a barrier against pathogenic microorganisms and toxins (37). Indeed, recent work has linked this type of diet with dysbiosis characterized by the proliferation of enterobacteria, including strains of adherent-invasive E. coli (AIEC) (38). This larger proportion of gut-resident Enterobacteriaceae may trigger susceptibility to infection by phages carrying the Stx gene (39), as these lysogenic strain derivatives can be a source of Stx. In addition, Enterobacteriaceae act as a source of LPS, favoring the development of a chronic, subclinical pro-inflammatory state which increases Stx-associated toxicity (40). Furthermore, high-fat, high-carbohydrate diets can stimulate the deleterious TLR4 inflammatory pathway by increasing LPS translocation and modulating brain functions (41, 42).

In addition to an adequate balance of macronutrients, especially fats and sugars, abundant research suggests that bioactive components present in food contribute to the proliferation of bacteria and promote the distribution of microbial communities, which can maintain homeostasis in the intestinal microenvironment and the brain-gut-microbiota axis (43).

## Conclusions

Thalamic and internal capsule cellular alterations correlated with neurological motor dysfunction and loss of sensitivity following unbalanced high-fats and high-carbohydrate diets. This work thus demonstrates that malnutrition should be considered a risk factor which may define disease severity following Stx2 intoxication.

## List of abbreviations

ANOVA: one-way analysis of variance; anti-GFAP: glial fibrillary acidic protein antibody; anti-Iba1: ionized calcium binding adaptor molecule 1 antibody; anti-NeuN: neuronal nuclei antibody; Gb3: globotriaosylceramide; HC: diet high in carbohydrates; HC+Stx2: diet high in carbohydrates treated with i.v Shiga toxin 2; HC+veh: diet high in carbohydrates treated with i.v vehicle; HC+HF: diet high in carbohydrates and high in fats; i.v: intravenous; HF: diet high in fats; HF+Stx2: diet high in fats treated with iv. Shiga toxin 2; HF+veh: diet high in fats treated with iv. vehicle; HC+HF+Stx2: diet high in carbohydrates and high in fats treated with i.v Shiga toxin 2; HC+HF+veh: diet high in carbohydrates and high in fats treated with i.v vehicle. HUS: hemolytic uremic syndrome; IOD: integrated optical density; anti-MBP: myelin basic protein antibody. ND: normal diet; ND+Stx2: normal diet treated with i.v Shiga toxin 2; ND+veh: normal diet treated with i.v vehicle; PBS: phosphate saline buffer; STEC: Stx producing *E. coli*; Stx2: Shiga toxin 2; VA-VL: ventral anterior and ventral lateral nuclei; veh: vehicle.

## Acknowledgments

We wish to thank German La Iacona for his invaluable technical advice and support in capturing fluorescent micrographs of this work.

## Funding

This work was supported by Agencia Nacional de PromociónCientífica y Tecnológica (ANPCyT) (PICT-2021-00656), Universidad de Buenos Aires (UBACyT) (20020190100186BA), and Consejo Nacional de InvestigacionesCientíficas y Técnicas (CONICET) (PIP 11220200101293CO), Argentina.

## Availability of data and materials

The datasets used and/or analyzed during the current study are available from the corresponding author on reasonable request.

## Authors’ contributions

Conceived of the experiment: JG. Conducted the experiment: DAM, NC, AC. Analyzed the data: DAM, NC, PAG, ELM, MDM, AP. Wrote the manuscript: AP, JG. All authors read and approved the final manuscript.

## Ethics approval

Experimental protocols and euthanasia procedures were reviewed and approved by the Institutional Animal Care and Use Committee of the School of Medicine at Universidad de Buenos Aires, Argentina (Resolution N° 3020/2019). All the procedures were performed in accordance with the EEOC guidelines for the care and use of experimental animals (EEC Council 86/609).

## Consent for publication

N/A

## Competing interests

The authors declare that they have no competing interests.

## Notes

### Competing Interest Statement

The authors have declared no competing interest.

